# First report of stop codon reassignment to tryptophan in members of the bacterial phylum *Actinomycetota*

**DOI:** 10.64898/2025.12.01.691617

**Authors:** Donovan H. Parks, Pierre-Alain Chaumeil, Maria Chuvochina, Philip Hugenholtz

## Abstract

Reassignment of stop codons is a significant evolutionary event with recoding of UGA to tryptophan being previously identified in only three bacterial phyla, the *Bacillota, Pseudomonadota*, and *Verrucomicrobiota*. Here, we present genomic evidence of this recoding in a fourth bacterial phylum, the *Actinomycetota*, specifically in the family *Eggerthellaceae*. We identify the UGA stop-to-tryptophan reassignment in 34 metagenome-assembled genomes recovered from the stool samples of diverse mammalian hosts, including equids and primates. Canonical markers for this reassignment are consistently observed including conserved UGA codons aligning to tryptophan, loss of release factor 2 (*prfB*), and presence of a tRNA^Trp^(UCA) gene. We infer that this reassignment occurred at least twice as the lineages containing recoded genomes are paraphyletic, forming two distinct groups separated by a third lineage with strains that continue to use UGA as a stop codon. These lineages represent three new *Eggerthellaceae* genera for which we propose the type species *Equivita altericodex, Gorillivita intestinalis*, and *Tapirivita inops* reflecting isolation source and genomic properties. Genomes representing these genera, including the non-recoded *Tapirivita* lineage, have reduced genomes and complete or partial loss of biosynthetic pathways, suggesting a transition to obligate symbiosis. Increasing host dependency may have facilitated stop codon reassignment in *Equivita* and *Gorillivita* species. This work expands the known phylogenetic diversity of UGA stop-to-tryptophan reassignment in the bacterial domain and establishes the *Eggerthellaceae* as a new focal point for understanding the evolutionary drivers of genetic code plasticity.

## Introduction

All organisms rely on a genetic code that translates nucleotide triplets (codons) on messenger RNA into specific amino acids or stop signals during protein synthesis [1]. Although this code is highly conserved, multiple independent reassignments have evolved in both cellular and organellar lineages [2]. Notably, several of these variations involve reassignment of stop codons to amino acids, a change identified in eukaryotic nuclear genomes [3, 4], mitochondria [5], and bacteria [6].

In bacteria, more than 99% of known species use the standard bacterial genetic code (code 11^1^). Two variants of this code are known, both involving reassignment of the UGA stop codon to either tryptophan (code 4) or glycine (code 25). Reassignment of UGA to tryptophan has been identified in obligate intracellular endosymbionts belonging to three bacterial phyla: the *Bacillota* order *Mycoplasmatales* mainly comprising endosymbionts of vertebrates, arthropods, and plants [7, 8]; *Pseudomonadota* insect endosymbionts *Hodgkinia cicadicola* [9], *Zinderia insecticola* [10], *Nasuia deltocephalinicola* [11] and *Stammera capleta* [12], and the dinoflagellate parasite *Fastidiosibacteraceae* species XS4 [13]; and two *Verrucomicrobiota* ciliate endosymbionts *Organicella extenuate* [14] and *Pinguicoccus supinus* [15]. Reassignment of UGA to glycine has been observed in all known species of *Candidatus* Absconditabacteria [16] and *Candidatus* Gracilibacteria [17], with both groups inferred to be symbionts based on lack of biosynthetic pathways for key metabolites [18, 19].

Bacterial genomes with UGA stop codon reassignment have several genomic features in common. This includes AT or GC compositional bias [9, 20] and reduced genome size (<2Mb) [2], with the insect and dinoflagellate endosymbionts having highly reduced genomes (<500 kb) [10, 13]. However, the key molecular signature of a recoded UGA stop codon is the loss of translational release factor RF2 and the mutation of the tryptophan or glycine tRNA gene to recognize the UGA codon [9, 16]. In the standard bacterial code, the three stop codons (UAA, UAG, and UGA) are recognized by translational release factors RF1 and RF2, with RF1 recognizing UAA and UAG and RF2 recognizing UAA and UGA [21]. Because both RF1 and RF2 recognize UAA, organisms that have reassigned UGA generally no longer require RF2 and thus typically lack the *prfB* gene that encodes RF2 [22]. Translation of UGA as an amino acid canonically requires a tRNA with the UCA anticodon. Accordingly, most organisms using genetic code 4 or 25 have a tRNA^Trp^(UCA) or tRNA^Gly^(UCA) gene, respectively. However, not all species with a UGA-to-tryptophan recoding have a tRNA^Trp^(UCA) gene. It has been experimentally demonstrated using the parasitic protist *Blastocrithidia nonstop* that a mutation of the anticodon stem of tRNA^Trp^(CCA) from 5 to 4 bp enables efficient translation of UGA [23]. This 4-bp anticodon stem tRNA^Trp^(CCA) gene was subsequently identified in the *Fastidiosibacteraceae* bacterium XS4, which uses genetic code 4 but lacks tRNA^Trp^(UCA) [13]. The functional versatility of UGA is further underscored by its role in encoding selenocysteine - the 21st amino acid - in a subset of organisms across all domains of life [24].

The phylogenetic distribution of UGA stop recodings across the bacterial domain remains incompletely mapped. Here, we provide genomic and phylogenetic evidence that the UGA codon has been reassigned to tryptophan in the *Actinomycetota* family *Eggerthellaceae*, a group of organisms commonly found in the gastrointestinal tract of humans and animals [25]. By analyzing a broad range of *Eggerthellaceae* genomes, we show that UGA recoding has occurred at least twice in this family accompanied by loss of multiple biosynthetic pathways, which we hypothesize indicates transition to an obligate host-associated lifestyle. This finding broadens the known taxonomic range of UGA recoding and establishes a new bacterial family for exploring the mechanisms and evolutionary consequences of genetic code evolution.

## Results

### Recovery of *Eggerthellaceae* MAGs

The reassignment of the UGA stop codon to tryptophan in two as-yet-uncultured *Eggerthellaceae* genera, JAUNQF01 and CAVGFB01, was identified as a result of applying the genetic code prediction tools Codetta [6] and gTranslate [26] to genomes in GTDB R10-RS226 [27]. To recover additional *Eggerthellaceae* genomes with this recoding, Sandpiper [28] was used to identify metagenomic samples in the Sequence Read Archive (SRA) [29] with evidence for the presence of JAUNQF01 (407 samples) or CAVGFB01 (31 samples) organisms (**Supp. Table 1**). Assembly and binning of these samples resulted in the recovery of 35 metagenome-assembled genomes (MAGs) estimated to be ≥70% complete and ≤10% contaminated, with all but three (MGR1, MGR3, and MGR6) predicted to use genetic code 4 (**Table 1**; **Supp. Table 2**).

**Table 1.** Genomic properties of *Eggerthellaceae* MAGs belonging to the genera JAUNQF01 and CAVGFB01.

| MAG ID | Genus | Species Cluster | Genetic code | Genome size (bp) | GC (%) | No. contigs | 16S rRNA | No. tRNA | Comp.# (%) | Cont.# (%) | Reference |
| --- | --- | --- | --- | --- | --- | --- | --- | --- | --- | --- | --- |
| GCA_030536315.1 | JAUNQF01 | 8 | 4 | 1,089,702 | 39.2 | 42 |  | 20 | 79.97 | 0.09 | [31] |
| GCA_030537355.1 | JAUNQF01 | 2 | 11 | 1,514,811 | 35.1 | 21 | partial | 20 | 93.05 | 0.22 | [31] |
| GCA_963628375.1 | CAVGFB01 | 1 | 4 | 1,044,520 | 32.0 | 158 | partial | 17 | 88.2 | 0.81 | [32] |
| MGR1 | JAUNQF01 | 10 | 11 | 1,445,008 | 35.1 | 15 |  | 20 | 97.94 | 0.16 | This Study |
| MGR2 | CAVGFB01 | 1 | 4 | 999,203 | 31.8 | 51 |  | 18 | 92.27 | 0.0 | This Study |
| MGR3 | JAUNQF01 | 2 | 11 | 1,524,775 | 35.1 | 21 | partial | 20 | 93.05 | 0.22 | This Study |
| MGR4 | JAUNQF01 | 4 | 4 | 1,400,689 | 37.8 | 119 | full | 18 | 92.34 | 0.09 | This Study |
| MGR5 | JAUNQF01 | 4 | 4 | 1,404,037 | 37.8 | 120 | full | 18 | 92.45 | 0.12 | This Study |
| MGR6 | JAUNQF01 | 2 | 11 | 1,523,567 | 35.1 | 23 | full | 20 | 93.05 | 0.24 | This Study |
| MGR7 | JAUNQF01 | 11 | 4 | 1,456,772 | 36.1 | 34 | full | 20 | 90.46 | 0.06 | This Study |
| MGR8 | CAVGFB01 | 1 | 4 | 1,220,050 | 32.1 | 91 |  | 15 | 91.5 | 0.46 | This Study |
| MGR9 | JAUNQF01 | 12 | 4 | 1,378,856 | 36.1 | 33 | partial | 20 | 89.43 | 0.05 | This Study |
| MGR10 | JAUNQF01 | 8 | 4 | 1,246,223 | 39.0 | 42 |  | 18 | 88.82 | 0.04 | This Study |
| MGR11 | JAUNQF01 | 7 | 4 | 1,605,346 | 37.5 | 20 |  | 20 | 91.61 | 1.21 | This Study |
| MGR12 | JAUNQF01 | 7 | 4 | 1,391,096 | 38.0 | 38 |  | 19 | 85.56 | 0.21 | This Study |
| MGR13 | JAUNQF01 | 5 | 4 | 1,007,078 | 39.9 | 144 | partial | 19 | 84.99 | 0.21 | This Study |
| MGR14 | CAVGFB01 | 1 | 4 | 1,429,972 | 31.3 | 81 |  | 19 | 95.92 | 2.79 | This Study |
| MGR15 | JAUNQF01 | 13 | 4 | 1,312,880 | 36.4 | 42 | partial | 17 | 84.6 | 0.68 | This Study |
| MGR16 | JAUNQF01 | 14 | 4 | 1,599,280 | 37.1 | 32 |  | 18 | 80.74 | 0.14 | This Study |
| MGR17 | JAUNQF01 | 6 | 4 | 985,589 | 39.1 | 59 |  | 19 | 86.23 | 1.49 | This Study |
| MGR18 | JAUNQF01 | 9 | 4 | 1,042,509 | 39.1 | 153 |  | 18 | 80.71 | 0.45 | This Study |
| MGR19 | JAUNQF01 | 15 | 4 | 1,123,380 | 33.3 | 57 |  | 15 | 79.38 | 0.34 | This Study |
| MGR20 | JAUNQF01 | 3 | 4 | 1,347,907 | 37.4 | 182 |  | 18 | 81.49 | 1.05 | This Study |
| MGR21 | CAVGFB01 | 1 | 4 | 814,140 | 31.7 | 70 |  | 19 | 78.05 | 0.52 | This Study |
| MGR22 | JAUNQF01 | 16 | 4 | 1,217,471 | 38.8 | 80 |  | 19 | 77.03 | 0.42 | This Study |
| MGR23 | JAUNQF01 | 4 | 4 | 1,576,414 | 37.6 | 76 |  | 18 | 93.84 | 4.02 | This Study |
| MGR24 | JAUNQF01 | 3 | 4 | 1,433,483 | 37.0 | 138 |  | 15 | 75.19 | 0.32 | This Study |
| MGR25 | JAUNQF01 | 6 | 4 | 939,565 | 38.9 | 151 |  | 18 | 73.82 | 0.15 | This Study |
| MGR26 | CAVGFB01 | 17 | 4 | 1,491,620 | 39.4 | 281 |  | 18 | 74.17 | 0.46 | This Study |
| MGR27 | JAUNQF01 | 8 | 4 | 1,333,865 | 39.1 | 66 |  | 20 | 88.84 | 3.43 | This Study |
| MGR28 | JAUNQF01 | 18 | 4 | 1,849,454 | 32.1 | 81 | partial | 18 | 89.26 | 3.63 | This Study |
| MGR29 | JAUNQF01 | 19 | 4 | 1,040,552 | 39.6 | 70 |  | 17 | 74.68 | 1.06 | This Study |
| MGR30 | JAUNQF01 | 9 | 4 | 1,085,064 | 39.0 | 93 |  | 19 | 77.23 | 2.55 | This Study |
| MGR31 | JAUNQF01 | 20 | 4 | 1,921,269 | 34.3 | 84 |  | 16 | 89.61 | 5.67 | This Study |
| MGR32 | JAUNQF01 | 21 | 4 | 1,170,523 | 38.9 | 126 |  | 18 | 72.52 | 2.52 | This Study |
| MGR33 | JAUNQF01 | 5 | 4 | 1,142,635 | 39.0 | 110 |  | 15 | 81.83 | 4.48 | This Study |
| MGR34 | JAUNQF01 | 22 | 4 | 1,202,115 | 37.5 | 97 |  | 17 | 76.42 | 5.04 | This Study |
| MGR35 | Novel | 23 | 4 | 1,065,035 | 37.1 | 199 |  | 16 | 74.03 | 4.57 | This Study |

The 35 recovered *Eggerthellaceae* MAGs were classified using GTDB-Tk [30] with 29 and 5 assigned to JAUNQF01 and CAVGFB01, respectively, and 1 comprising a new genus in this family (**Figure 1**; **Table 1**). These 35 MAGs, together with a single CAVGFB01 and 2 JAUNQF01 MAGs in GTDB R10-RS226, were grouped into 23 species clusters using a 95% average nucleotide identity (ANI) criterion. Nine of the species clusters contained multiple MAGs, with only cluster sp5 (comprised of MGR13 and MGR33) not being resolved as monophyletic (**Figure 1**; **Table 1**). We refer to this set of 38 MAGs as the Mammalian Gut Recoding (MGR) clade and to the paraphyletic subsets of MAGs predicted to use genetic code 4 as MGR-GC4.

**Figure 1.**
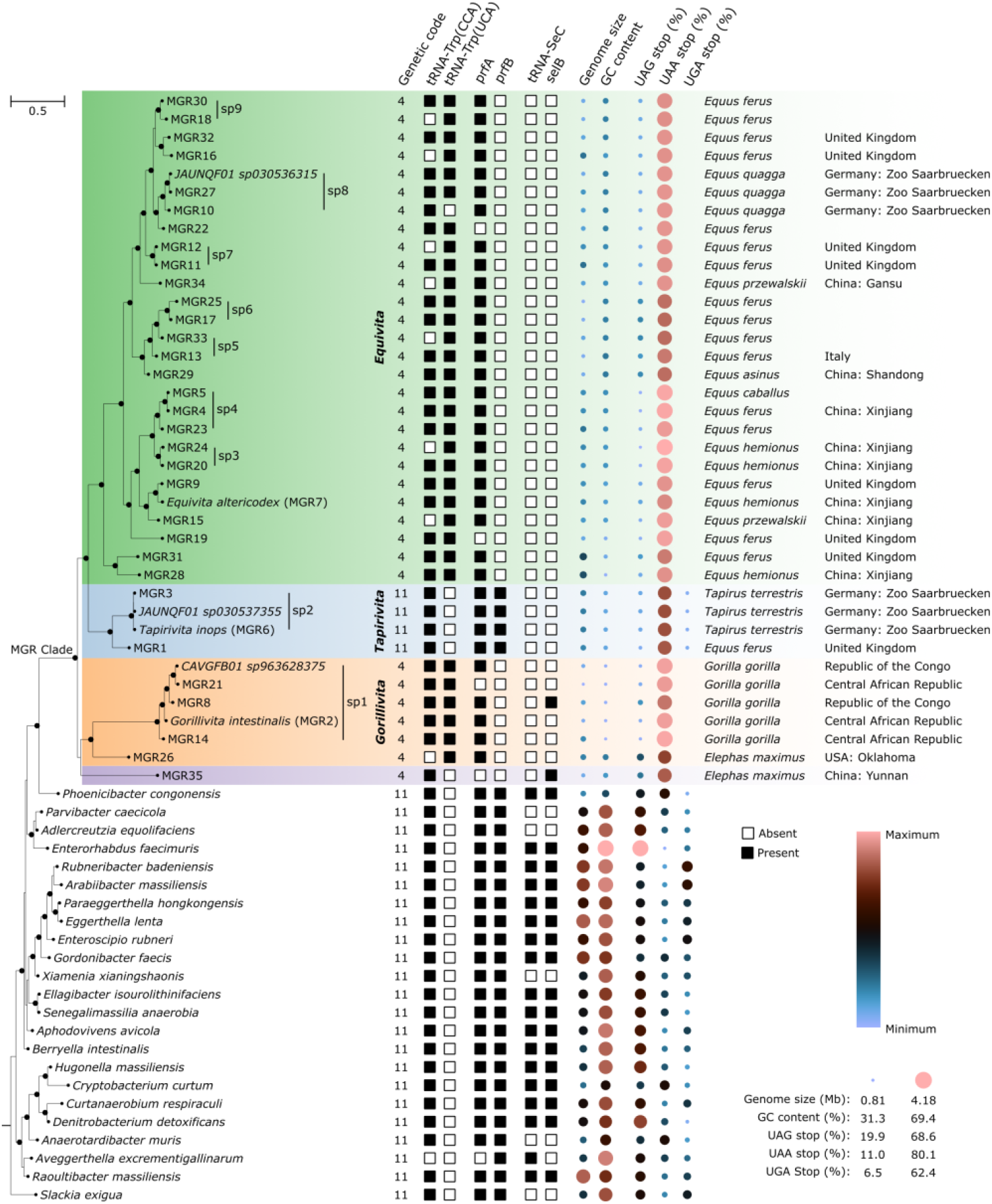
Maximum-likelihood phylogenetic tree inferred from alignment of 120 bacterial marker genes with IQ-Tree and the LG+F+C40 evolutionary model. *Equivita* MAGs (green shading), *Tapirivita* MAGs (blue shading), *Gorillivita* MAGs (orange shading) and a single recoded MAG belong to a new genus (purple shading) are shown along with isolate genomes from *Eggerthellaceae* genera. Genomes belonging to the same species are denoted by black horizontal lines. The tree was rooted on the UBA8131 species *WRGR01 sp009787085* (GCA_009787085.1) as this family is consistently found to be sister to *Eggerthellaceae*. Black dots on internal branches indicate strong support with UFBoot ≥95% and SH-aLRT ≥80%. Genomic properties that indicate a UGA stop to tryptophan recoding are shown for each genome. Minimum and maximum values for genome size, %GC content, and stop codon usage were calculated over all MAGs in the figure and isolate genomes from 61 *Eggerthellaceae* species in GTDB R10-RS226. *prfA* = translational release factor RF1; *prfB* = translational release factor RF2; *selB* = selenocysteine-specific elongation factor.

### MGR MAGs represent new genera and species within *Eggerthellaceae*

Here we propose type genomes and names for MGR taxa following the SeqCode [34], a sequence-based nomenclatural code. We propose that the JAUNQF01 placeholder genus be divided into two genera, one comprising the MAGs using genetic code 11 and the other the MAGs using genetic code 4 (**Figure 1**). The proposal for two genera is supported by relative evolutionary divergence [35] and percent identity of 16S rRNA gene sequences [36] from these two clades (*see Supp. Note 1*). We designate MGR6 as the type genome for the GC11 JAUNQF01 sequences and propose the name *Tapirivita inops* gen. nov., sp. nov. to reflect the host species of this MAG and its reduced genome size. For the GC4 JAUNQF01 sequences, we designate MGR7 as the type genome and propose the name *Equivita altericodex* gen. nov., sp. nov. to reflect the host species of this MAG and recoding of the UGA stop codon. Finally, we designate MGR2 as the type genome for the CAVGFB01 sequences and propose the name *Gorillivita intestinalis* gen. nov., sp. nov. to reflect the host species of this MAG and the MGR MAGs being associated with the gastrointestinal tract of mammals. Protologues for new taxa are provided in **Supp. Table 3**. The MAGs selected as type genomes meet most criteria required to be considered high quality [37], as they have an estimated completeness >90% with minimal estimated contamination, are comprised of ≤51 contigs, and have ≥18 tRNA genes (**Supp. Table 2**). However, MAGs often lack the rRNA operon and this is reflected in the selected MAGs with only MGR6 containing the 5S, 16S, 23S rRNA genes, MGR7 having only a 16S rRNA gene, and MGR2 missing all three of these rRNA genes.

### Phylogenetic relationship of *Eggerthellaceae* genomes

A maximum-likelihood tree was inferred to examine phylogenetic relationships among *Eggerthellaceae* genomes and the 38 MGR MAGs (**Figure 1**; **Supp. Table 1**). Notably, the *Equivita* and *Gorillivita* MAGs form two distinct lineages that use genetic code 4, indicating either two independent recoding events or a single recoding event followed by a reversion to genetic code 11 in the *Tapirivita* lineage. To confirm these evolutionary relationships, 24 phylogenetic trees were inferred using varying marker sets, alignments, inference methods, and evolutionary models (**Supp. Table 4**). All trees indicate that the MGR clade comprising *Equivita, Gorillivita, Tapirivita*, and MGR35 is monophyletic (**Figure 1**; **Supp. Figure 1**). Notably, the GC4 genera *Equivita* and *Gorillivita* were consistently paraphyletic and separated by the GC11 genus *Tapirivita*. GTDB-Tk classified MGR35 as a new *Eggerthellaceae* genus consistent with its unstable phylogenetic placement: 14 of 24 trees placed it sister to *Gorillivita*, nine placed it as the most basal lineage in the MGR clade, and one placed it between *Gorillivita* and *Tapirivita* (**Supp. Figure 1**). Most trees (20 of 24) place the MGR clade as a non-basal lineage within the *Eggerthellaceae* except for four trees inferred with a more heuristic inference method (FastTree) and simple evolutionary models (WAG or LG+G; **Supp. Figure 1**).

### Genomic properties supporting recoding

UGA-to-tryptophan recoding was predicted by two complementary tools: gTranslate, a machine learning method for inferring the genetic code used by prokaryotic genomes, and Codetta, a homology-based approach that predicts recodings by examining alignments of conserved proteins. Across the 34 MGR-GC4 MAGs (i.e. *Equivita* MAGs, *Gorillivita* MAGs, and MGR35), Codetta identified 249 to 823 (mean = 488) instances where UGA aligned to a conserved tryptophan residue (**Supp. Table 2**). No additional deviations from the standard bacterial genetic code were identified by Codetta. We identified five proteins that illustrate the alignment of UGA codons to conserved tryptophan residues across the 34 MGR-GC4 MAGs contrasted to the 4 GC11 *Tapirivita* MAGs and 3 outgroup *Eggerthellaceae* genomes (**Figure 2**).

**Figure 2.**
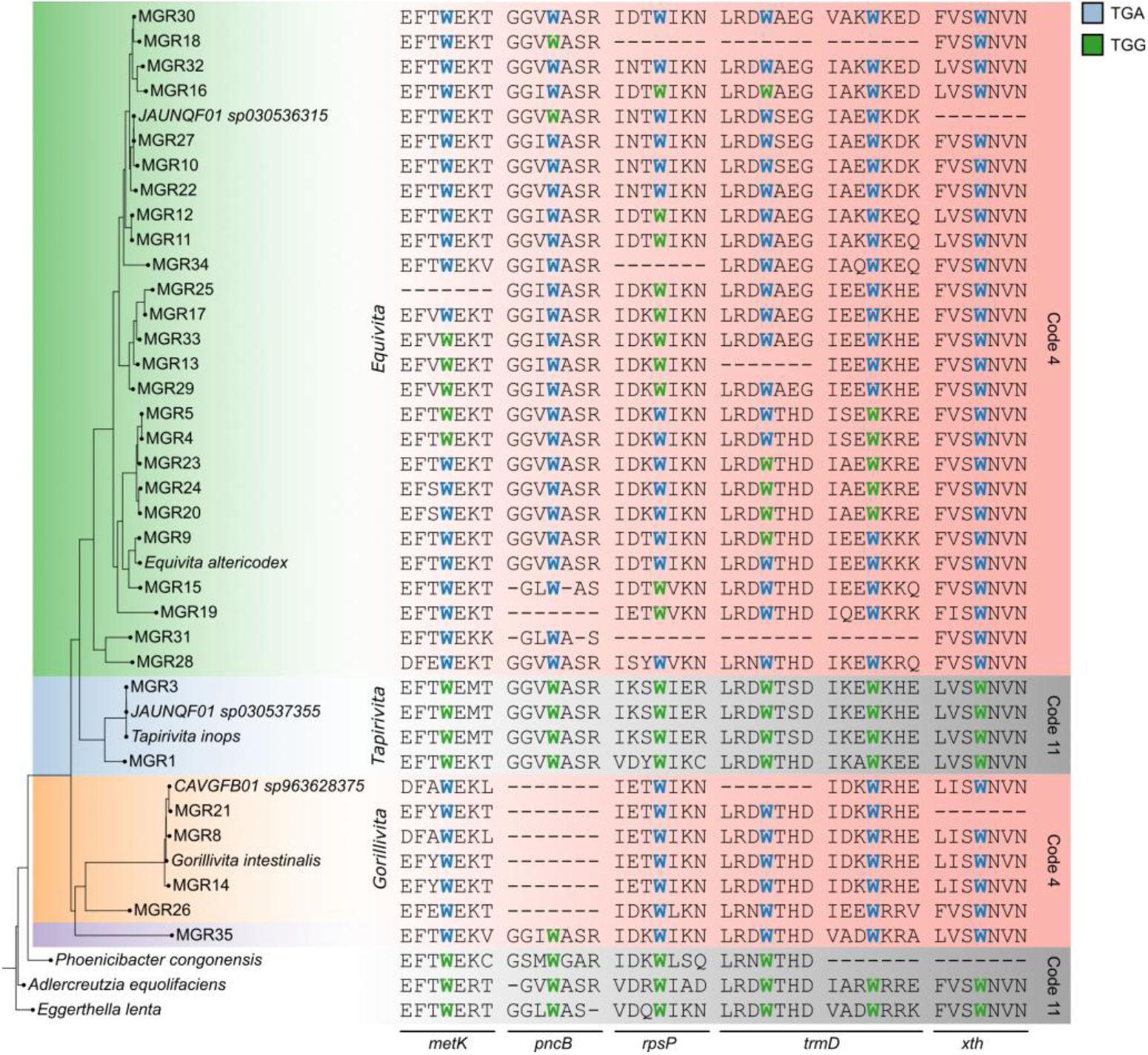
Aligned proteins indicating the TGA codon codes for tryptophan in the 27 *Equivita* MAGs, 6 *Gorillivita* MAGs, and the MGR35 MAG. Conserved tryptophan (W) residues are encoded by TGA (blue) or TGG (green). Partial sequences are for proteins encoded for by the genes *metK* (methionine adenosyltransferase), *pncB* (nicotinate phosphoribosyltransferase), *rpsP* (ribosomal protein S16), trmD (guanine(37)-N(1))-methyltransferase) and *xth* (exodeoxyribonuclease III).

The MGR-GC4 MAGs exhibit genomic properties that are typical of stop codon recoded organisms (**Table 2**). This includes having a relatively small genome size (mean = 1.28 Mb) and low %GC content (mean = 36.8%) in comparison to other *Eggerthellaceae* species (mean genome size = 2.23 Mb; mean GC = 58.6%; **Supp. Table 1**). All MGR MAGs have a homolog of *prfA* (RF1; **Figure 1**), except for four that we attribute to these MAGs being incomplete (mean estimated completeness of 85.0% ± 7.4%; **Supp. Table 2**). By contrast, all MGR-GC4 MAGs lack a homolog of *prfB* (RF2; **Figure 1**) and sequence similarity searches against *Eggerthellaceae* RF2 proteins indicated the absence of even pseudogenized *prfB* sequences in these MAGs (*see Methods*). Loss of *prfB* is common in genomes using genetic code 4, as this gene is dispensable when UGA no longer serves as a stop codon (**Table 2**) [7, 38]. Additionally, the majority of MGR-GC4 MAGs contain tRNA^Trp^ genes with both CCA and UCA anticodons (**Figure 1**), the latter enabling recognition of UGA as tryptophan [39]. Presence of a tRNA^Trp^(UCA) gene is common among bacterial species using genetic code 4 (**Table 2**), though not ubiquitous as exemplified by the *Fastidiosibacteraceae* XS4 bacterium [13].

**Table 2.** Genomic properties of select bacterial species that use genetic code 4.

|  | *Equivita* and<br>*Gorillivita* | *Mycoplasmatales*<br>*spp.* | *Hodgkinia*<br>*cicadicola* | *Nasuia*<br>*deltocephalinicola* | *Stammera*<br>*capleta* | *Zinderia*<br>*insecticola* | *Pinguicoccus*<br>*supinus* |
| --- | --- | --- | --- | --- | --- | --- | --- |
| No. genomes | 34 | 220 | 13 | 24 | 49 | 1 | 1 |
| No. species | 21 | 160 | 1 | 1 | 1 | 1 | 1 |
| %GC content | 31.3 to 39.9<br>mean = 36.8 | 22.4 to 40.0<br>mean = 27.1 | 38.7 to 58.4<br>mean = 47.6 | 15.2 to 18.0<br>mean = 16.9 | 13.9 to 20.4<br>mean = 16.9 | 13.5 | 25.1 |
| Genome size<br>(Mb) | 0.81 to 1.92<br>mean = 1.28 | 1.18 to 2.23<br>mean = 1.34 | 0.11 to 0.15<br>mean = 0.14 | 0.11 to 0.14<br>mean = 0.11 | 0.22 to 0.34<br>mean = 0.29 | 0.21 | 0.16 |
| tRNA<sup>Trp</sup>(CCA) | 27 of 34 | 211 of 220 | No | No | Yes | Yes | No |
| tRNA<sup>Trp</sup>(UCA) | 32 of 34 | 219 of 220 | Yes | Yes | No | No | Yes |
| *prfB* (RF2) | No | No | No | No | 15 of 49 | No | No |
| tRNA-SeC | No | No | No | No | No | No | No |
| *selB* | 1 of 34 | 2 of 220 | No | No | No | No | No |
| UAA stop (%) | 65.2 to 80.1<br>mean = 75.6 | 65.5 to 91.3<br>mean = 80.8 | 30.5 to 68.8<br>mean = 55.0 | 87.1 to 96.6<br>mean = 91.8 | 88.2 to 97<br>mean = 92.8 | 95.8 | 78.5 |
| Host | Mammalian gut | Vertebrates,<br>arthropods,<br>plants | Cicadas<br>(*Cicadoidea*) | Leafhoppers<br>(*Hemiptera*) | Leaf beetles<br>(*Cassidinae*) | Spittlebugs<br>(*Cercopoidea*) | Ciliate<br>(*Euplotes*<br>*vanleeuwenhoekii*) |
| Phylum | *Actinomyetota* | *Bacillota* | *Pseudomonadota* | *Pseudomonadota* | *Pseudomonadota* | *Pseudomonadota* | *Verrucomicrobiota* |
| Reference | This Study | [7] | [9] | [11] | [12] | [10] | [15] |

MGR-GC4 MAGs preferentially use the UAA instead of UAG stop codon (mean frequency of 75.9% vs 24.1%; **Figure 1**; **Supp. Table 2**) presumably reflecting their low %GC content [40]. Notably, the GC11 *Tapirivita* MAGs also favour the UAA stop codon (mean = 67.5%; minimum = 66.7%) and rarely use the UGA stop codon (mean = 9.2%; maximum = 9.6%). This contrasts with other *Eggerthellaceae* species, where usage is more even across the 3 stop codons (UAG = 44.6%, UAA = 34.9%, and UGA = 20.5%; **Supp. Figure 2**). The highest usage of UAA is 61.6%, and only 15 of 212 (7.1%) genomes exhibit UGA usage at or below 9.6% (**Supp. Table 2)**. This suggests that the *Tapirivita* MAGs have experienced similar selective pressures as the MGR-GC4 MAGs that resulted in reduced %GC content, preferential usage of the UAA stop codon, and reassignment of UGA to tryptophan.

Collectively, these results support UGA encoding tryptophan in members of the MGR clade, with the notable exception of the two *Tapirivita* species where genomic evidence indicates they use UGA as a stop codon.

### Evidence for multiple recoding events

Loss of *prfB* (RF2) has been proposed as the common force driving UGA stop recoding events [9]. The loss of this gene is nearly universal in organisms that have recoded the UGA stop codon as the presence of RF2 would result in early termination of proteins at UGA codons (**Table 2**) [22]. The phylogenetic relationship of MAGs in the MGR clade suggests two independent recoding events or a single recoding event followed by reversion to code 11 in the genus *Tapirivita* (**Figure 1**). Examination of the gene neighbourhood around *prfB* (RF2) and the phylogeny of this gene support two independent recoding events. The *secA* and *prfB* genes have previously been reported to constitute an operon in *Bacillus subtilis* [41] and we observed that these genes are adjacent or separated by a single gene in all *Eggerthellaceae* species using genetic code 11 (**Figure 3a**; **Supp. Table 2**). The conserved adjacency of *secA* and *prfB* in the *Tapirivita* MAGs suggests *prfB* was not lost in this lineage, as re-acquisition via horizontal gene transfer into this specific locus is unlikely. The *prfB* phylogeny further supports that this gene has not been lost by *Tapirivita* MAGs, as it is largely congruent with the species tree—notably, *P. congonesis* is the closest genetic code 11 species to *Tapirivita* MAGs in both trees (**Figures 1** and **3b**). The implications of multiple recoding events in this lineage are discussed in the next section.

**Figure 3.**
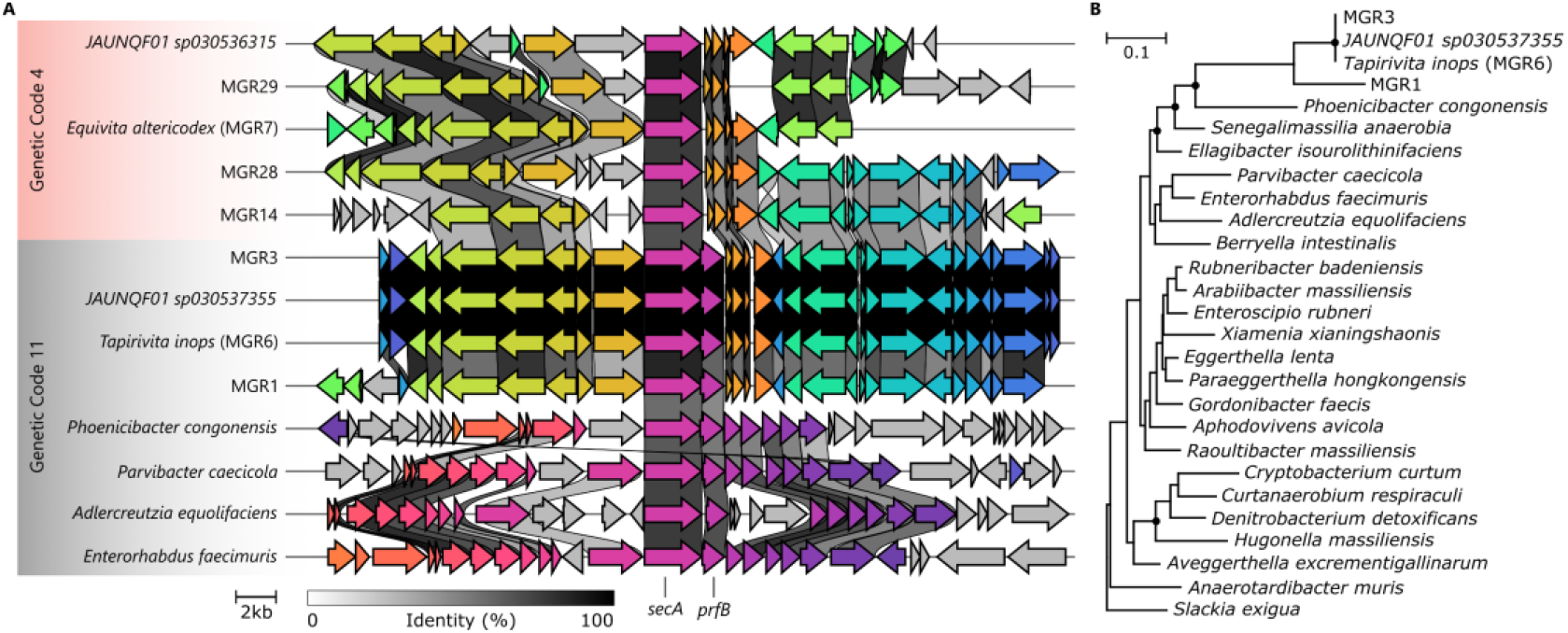
Gene neighbourhood around *prfB* (**A**) and the phylogeny of this gene (**B**) support two independent recoding events. **(A)** Gene neighbourhood around *secA* illustrating its close proximity to *prfB* (RF2) in *Eggerthellaceae* genomes using genetic code 11 and the loss of *prfB* in MGR-GC4 MAGs. Genes in adjacent genomes with >35% identity are connected by a grey ribbon. **(B)** Maximum-likelihood phylogenetic tree of *prfB* inferred with IQ-Tree and the Q.pfam+I+G4 evolutionary model. The tree was rooted on *Slackia exigua* as this species was basal in the *Eggerthellaceae* species tree (Figure 1). Black dots on internal branches indicate strong support with UFBoot ≥95% and SH-aLRT ≥80%. *secA* = protein translocase subunit.

### Inferred transition of MGR organisms to obligate host dependency

Stop codon reassignment is often associated with a transition to obligate host association as seen for *Mycoplasmatales* and *Pseudomonadota* endosymbionts [42]. Such transitions are marked by reductive evolution that typically results in reduced genome size and change in %GC content, as already noted for MGR genomes (**Table 1**), and loss of biosynthetic pathways and cell envelope structures [42]. Loss of biosynthetic capability is linked to availability and increasing reliance on host metabolites and deterioration of the cell envelope has been linked to osmotic homeostasis in a eukaryotic host cell and reduced immune triggering [43]. All MGR genomes (including the GC11 *Tapirivita* species) displayed complete or partial loss of biosynthetic pathways relative to other *Eggerthellaceae* genomes, including for amino acids (histidine, leucine, isoleucine, threonine, proline, lysine and arginine), vitamins (B6, B9), and core metabolism (fatty acid synthesis, TCA cycle; **Supp. Table 5**). By contrast, genes for synthesis and maintenance of the cell envelope remain mostly intact (**Supp. Table 6**), which has been observed in GC11 obligate insect endosymbionts including *Buchnera, Blochmanniella*, and *Wigglesworthia* [44–46] and appears to be the case in at least one GC4 symbiont, *Stammera capleta* [12]. Together, these genomic features suggest that MGR clade species have undergone genome reduction and transitioned to a host-dependent lifestyle in the mammalian gut.

The presence of the GC11 genus *Tapirivita*, with hallmark features of obligate symbionts, in the midst of GC4 species in the MGR clade (**Figure 1**) calls into question the timing of UGA reassignment. UGA to tryptophan recoding is unlikely to be reversible [47, 48] suggesting that at least two independent recoding events have occurred in the MGR lineage (*see discussion above*) in bacteria that had already undergone genome reduction. Therefore, stop codon recoding is more likely to be a consequence of genome reduction and host adaptation rather than a cause. Low usage of UGA stop codons in the two *Tapirivita* species (∼9%; **Figure 1**) suggests that they may be primed for a recoding event and thus may offer compelling targets for studying this phenomenon.

### Host association and biogeography of MGR genera

Bacteria that use genetic code 4 are typically symbiotic organisms with reduced genomes that reside in a range of insect, arthropod, vertebrate, plant, and protist hosts (**Table 2**). MGR organisms follow this pattern and were exclusively recovered from mammalian stool samples. *Equivita* MAGs were exclusively associated with hosts in the *Perissodactyla* genus *Equus* (e.g., horses, zebras; **Figure 1**), but metagenomic profiling indicates a broader host range spanning the additional *Perissodactyla* species *Diceros bicornis* (black rhinoceros) along with *Ovis aries* (sheep), *Crocuta crocuta* (hyena), and *Elephas maximus* (elephants; **Figure 4**; **Supp. Table 1**). By contrast, *Tapirivita* species may be restricted to hosts in the order *Perissodactyla* with MAGs recovered from *Equus* and *Tapirus terrestris* (tapir) stool samples and metagenomic profiling only increasing the host range to include the *Perissodactyla* species *Rhinoceros unicornis* (Indian rhinoceros). *Gorillivita* MAGs were recovered from *Gorilla gorilla* and *Elephas maximus* fecal samples, and metagenomic data expanded this host range to include *Pan troglodytes* (Chimpanzee) and *Loxodonta africana* (African bush elephant). Notably, this suggests *Gorillivita* species may not be found in *Perissodactyla* hosts.

**Figure 4.**
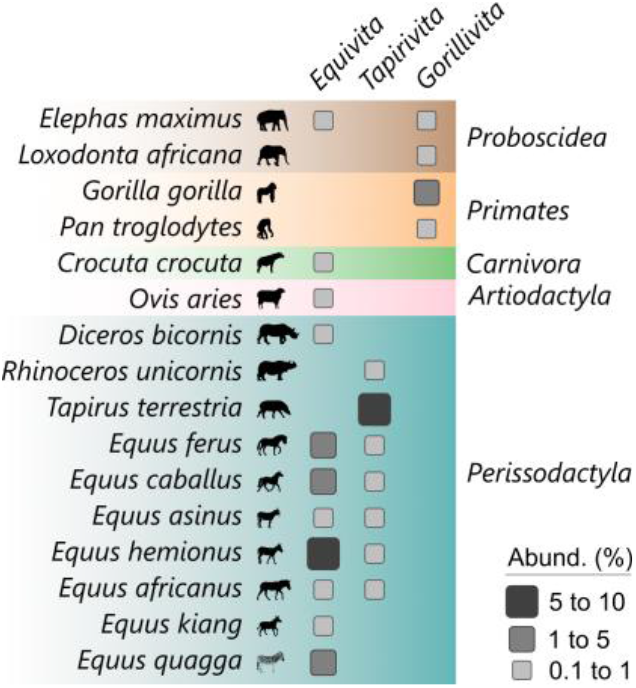
Hosts associated with the MGR genera *Equivita, Tapirivita*, and *Gorillivita* identified by metagenomic profiling. The maximum relative abundance of MGR genera across profiled samples is indicated for each host species using grey boxes. Host are organized and colored by taxonomic order.

*Equivita* and *Tapirivita* are widely distributed due to the geographic range of their mammalian hosts. Equine stool samples containing these MGR genera were collected in Europe (e.g., United Kingdom, Italy, Denmark), Asia (i.e., China, Iran), and North America (i.e., Canada; **Supp. Table 1**). The biogeography of *Equivita* also includes the South and East African countries of South Africa, Kenya, Tanzania due to its host association with *D. bicornis, C. crocuta*, and *E. quagga*, respectively. Stool samples from *T. terrestris* (tapir) were collected from the Saarbrücken Zoo in Germany but suggest that *Tapirivita* may be present in South America, the tapir’s native habitat. *Tapirivita* was also identified in a *R. unicornis* stool sample from Nepal. The geographic range of *Gorillivita* includes Central Africa (Congo, Central African Republic, Gabon) where it is associated with gorillas and/or chimpanzees (**Supp. Table 1**). *Gorillivita* was also identified in the stool samples of two captive elephant species, *E. maximus* and *L. africana*, which reside in the Oklahoma [49] and Beijing [50] zoos, respectively. This suggests a wider geographic distribution of *Gorillivita* species as the native range for *E. maximus* spans the Indian subcontinent and Southeast Asia while *L. africana* is found across sub-Saharan Africa.

## Discussion

Bacteria using genetic code 4 (GC4) have previously only been identified in the phyla *Bacillota* [51], *Pseudomonadota* [10], and *Verrucomicrobiota* [14, 15]. The MGR-GC4 genera *Equivita* and *Gorillivita* belong to the phylum *Actinomycetota* thereby expanding the number of phyla known to contain organisms using this recoding. The average genome size of MGR-GC4 MAGs (mean = 1.28 Mb) and GC4 *Bacillota* order *Mycoplasmatales* genomes (mean = 1.34 Mb) are similar (**Table 2**), and while small compared to most bacterial organisms (∼5 Mb) [52] are much larger than the highly reduced GC4 *Pseudomonadota* genomes (<0.5 Mb) of insect or dinoflagellate endosymbionts [10, 13]. These organisms often have extremely low %GC content (**Table 2**) as exemplified by the insect endosymbionts *Nasuia deltocephalinicola* (mean = 16.9%) [11], *Stammera capleta* (mean = 16.9%) [12], and *Zinderia insecticola* (13.5%) [10] and to a lesser extent the dinoflagellate endosymbiont XS4 (28.2%) [13]. The majority of *Mycoplasmatales* species also exhibit low %GC content (mean = 27.1%). However, *Hodgkinia cicadicola* with a %GC content as high as 58.4% [9] demonstrates that low %GC content is not a requirement of organisms that use genetic code 4 and the MGR-GC4 MAGs with %GC ranging from 31 to 40% (mean = 36.8%) support a wider %GC content range for GC4 organisms than previously appreciated.

While this study has primarily focused on evidence supporting a UGA recoding and establishing the phylogenetic relationship between *Eggerthellaceae* species, the presence of both GC4 and GC11 species in the MGR clade makes it an interesting system for investigating the mechanisms leading to and the results of UGA being reassigned to tryptophan. However, any such investigations will benefit from the recovery of additional *Tapirivita* MAGs as this genus currently comprises only 4 MAGs from 2 species, with the 3 *Tapirus terrestris* MAGs (MGR3, MGR5, JAUNQF01 sp030537355) being recovered from biological replicates of, presumably, a single tapir at the Saarbrücken Zoo in Germany [31]. The majority of *Eggerthellaceae* species outside the MGR clade have the genes required to incorporate selenocysteine into proteins using a translational recoding of UGA, *i*.*e*. a selenocysteine-specific tRNA (tRNA^SeC^) and a specialized translation factor (*selB*) that bring Sec-tRNA^SeC^ to the ribosome at UGA sites with a downstream SECIS element (**Figure 1**) [53, 54]. Consequently, the MGR clade being sister to these *Eggerthellaceae* species and lacking selenocysteine-associated genes suggests that translational recoding of UGA may have acted as a precursor to the MGR clade recoding UGA to tryptophan. To our knowledge, use of UGA for selenocysteine potentially priming UGA stop codon recoding has not been previously proposed, and the mechanisms that would support this change are unclear. One hypothesis is that selenocysteine incorporation competes with the RF2 release factor for the UGA codon [55, 56] resulting in a preference for alternative stop codons and a reduction in the use of the RF2, ultimately resulting in the loss of UGA as a stop codon and the need for the RF2 gene.

The MGR-GC4 MAGs are broadly consistent with both the ‘codon capture’ [20, 57] and ‘ambiguous translation’ [5, 58] models for the reassignment of the UGA stop codon to tryptophan. The codon capture model proposes that this reassignment occurs after the UGA stop codon has been driven to extinction because of reduced %GC content that favours a mutational bias to the UAA stop codon. The UGA codon would then be under no pressure to be maintained as a stop codon and could be re-introduced as recognizing tryptophan. The MGR-GC4 MAGs are consistent with this model as they have lower %GC content (mean = 36.8%) than other *Eggerthellaceae* species (mean GC = 58.6%). While %GC content of the extant MGR-GC4 MAGs is arguably not sufficiently low to support the codon capture model, it is possible that the ancestor(s) of these extant species had substantially lower %GC content that would be compatible with this model, or another mechanism not reliant on low %GC caused the recoding. The ambiguous translation model proposes that a single codon is temporarily read in two different ways, with a subsequent loss of the original meaning of the code. McCutcheon et al. [9] adapted this model to explain UGA recoding in the high %GC *H. cicadicola* where a mutated tRNA^Trp^(CCA) would allow recognition of both UGG and UGA. This would be followed by loss of RF2 as the genome mutates to using UAA and UAG as stop codons and UGA as tryptophan. For *H. cicadicola*, subsequent mutation of the tRNA^Trp^ anticodon would result in these genomes exclusively having tRNA^Trp^(UCA). This model is largely consistent with the MGR-GC4 MAGs that have also lost the RF2 gene and have a tRNA^Trp^(UCA) gene, though also retain a tRNA^Trp^(CCA) gene (**Figure 1**). This could be the result of an initial duplication of the tRNA^Trp^(CCA), which is plausible as multiple copies of this gene were identified in 14 *Eggerthellaceae* species (**Supp. Table 2**).

## Conclusions

We present genomic evidence for UGA stop-to-tryptophan recoding in 34 MAGs comprising 3 genera and 21 species in the family *Eggerthellaceae*. These MAGs were found exclusively in mammalian stool samples, exhibit both host and geographic structuring, and are the first members of the bacterial phylum *Actinomycetota* identified with this reassignment. They have features consistent with a transition to obligate host association but are notable for having moderate %GC content compared to the AT rich genomes typical of GC4 obligate symbionts, raising questions about the evolutionary mechanism that drove reassignment. We infer that UGA stop-to-tryptophan recoding occurred at least twice in the *Eggerthellaceae* based on phylogenetic and gene synteny evidence that suggests this transition was a consequence of genome reduction and host adaptation rather than a driver. Thus, the family *Eggerthellaceae* provides a new focus for furthering our understanding of the mechanisms leading to genetic code reassignments.

## Material and methods

### Recovery of additional *Eggerthellaceae* MAGs

Metagenomic samples from the SRA [29] likely to result in recovery of CAVGFB01 or JAUNQF01 MAGs were identified using Sandpiper 1.0.0 [28]. Specifically, we processed SRA runs where either genus was estimated to be at ≥1x coverage and ≥0.1% abundance with Aviary 0.12.0 (https://github.com/rhysnewell/aviary) using default parameters. This resulted in 438 samples (31 CAVGFB01; 407 JAUNQF01) being processed (**Supp. Table 1**). Recovered MAGs were classified using GTDB-Tk v2.4.1 [30] which resulted in the identification of 15 CAVGFB01 MAGs, 84 JAUNQF01 MAGs, and single MAG from a novel *Eggerthellaceae* genus predicted to use genetic code 4. CheckM2 1.1.0 [33] was used to estimate the quality of these 100 MAGs and the 35 MAGs (5 CAVGFB01, 29 JAUNQF01, and the novel *Eggerthellaceae* MAG) with ≥70% completeness, ≤10% contaminated, and a quality score ≥50 (defined as completeness - 5×contamination) used in subsequent analyses. These 35 MAGs came from 27 SRA runs which have a minimum estimated CAVGFB01 or JAUNQF01 coverage of 6.8x and abundance of 0.17%.

Aviary used the following programs to recover MAGs: fastp 1.0.1 [59], metaSPAdes 4.0.0 [60], CoverM 0.7.0 [61], minimap 2.18 [62], MetaBAT 2.15 [63], SemiBin2 2.2.0 [64], Vamb 5.0.4 [65], DAS Tool 1.1.2 [66], CheckM 1.2.4 [67], CheckM2 1.1.0 [33], and pplacer 1.1 [68]. The translation table of *Eggerthellaceae* genomes, including the 35 quality-filtered MGR MAGs, were predicted using gTranslate 1.0 [26] and Codetta 2.0 [6] using default settings.

### Assignment of MGR MAGs to species clusters

The 35 MGR MAGs along with the 2 JAUNQF01 and single CAVGFB01 MAGs in GTDB R10-RS226 were organized into operational species clusters using Galah 0.4.2 (github.com/wwood/galah). Galah uses the same representative selection criteria and clustering methodology as GTDB [69], except the presence of 16S rRNA genes is not considered.

### Inference of species trees

The phylogenetic relationship of *Eggerthellaceae* species was determined using genomes from GTDB R10-RS226 and the 35 MGR MAGs recovered in this study (**Supp. Table 2**). Specifically, trees were inferred using the 212 (of 297) GTDB species representatives within the *Eggerthellaceae* family with a CheckM2 completeness ≥80%, contamination ≤5%, and consists of ≤200 contigs along with the CAVGFB01 (GCA_963628375.1) and JAUNQF01 (GCA_030537355.1, GCA_030536315.1) genomes. The trees were rooted using the *WRGR01 sp009787085* genome GCA_009787085.1, the highest-quality MAG in the family UBA8131, which is sister to *Eggerthellaceae* according to the GTDB R10-RS226 reference tree.

Trees were inferred using four marker sets, varying methods for producing and trimming the multiple sequence alignment (MSA), and different tree inference methods and models of evolution (**Supp. Table 4**). The four marker sets considered were the 120 bacterial (bac120) marker genes used by GTDB [70], a set of 16 ribosomal proteins (rp1) [71], a set of 23 ribosomal proteins (rp2) [17], and a set of 339 core *Eggerthellaceae* genes. The bac120 genes were identified using GTDB-Tk 2.4.1 [30]. The rp1 and rp2 genes were identified using EasyCGTree 4.2 [72] run with default values. The core *Eggerthellaceae* gene set was established using a modified version of Panaroo 1.5.2 [73] that allowed the genetic code for each genome to be individually specified. Panaroo was run in strict mode using a sequence identify threshold of 60%, protein family identity threshold of 50%, length difference threshold of 60%, and core gene threshold of 70%. Genes were aligned using HMMER 3.4, MUSCLE 5.1 [74] run in ‘-align’ mode, or MAFFT 7.505 [75] run using the L-INS-i alignment method and --maxiterate set to 1000. Trees were inferred for each MSA without filtering, after filtering with TrimAl 1.4.1 [76] run with -automated1, and after further filtering with WitChi [77] run with the quartic pruning algorithm in ‘touchdown’ mode.

Maximum-likelihood trees were inferred with IQ-Tree 2.4.0 [78] with the evolutionary model selected using ModelFinder [79]. Support values were determined using 1000 replicates of both the SH approximate likelihood ratio test (SH-aLRT) [80] and ultrafast bootstraps (UFBoot) [81]. For the bac120 marker set, trees were also inferred using FastTree 2.1.11 [82] using i) the full set of GTDB species representatives, MSA filtering as implemented in GTDB-Tk 2.4.1, and the WAG evolutionary model in order to replicate the tree inference method used by the GTDB [69], and ii) the LG+G model with the MSA filtered using TrimAl and WitChi as described above in order to evaluate the impact of tree inference methods.

The tree shown in Figure 1 was inferred over the full set of 215 GTDB species representatives and MGR MAGs as described above but pruned to contain only 1 genome per genus with a Latin name. The genome for the type species of the genus was retained when available.

### Inference of *prfB* gene tree

*prfB* genes >180 bp were aligned with MAFFT 7.505 using the --auto argument. IQ-Tree 2.4.0 was used to infer the gene tree with ModelFinder used to select the most appropriate evolutionary model and support values determined using 1000 replicates of SH-aLRT and UFBoot.

### Identification of genes

MGR MAGs along with additional *Eggerthellaceae* genomes were annotated with Prokka 1.14.6 [83] using the bacterial reference databases. Aragorn 1.2.38 [84] was used internally by Prokka to identify tRNA genes. Prokka annotations were inspected to identify tRNA^Trp^ gene, peptide chain release factor 1 (RF1, *prfA*) and 2 (RF2, *prfB*) genes, genes comprising the *selABCD* operon, and genes comprising the 16S-23S-5S rRNA operon. The lack of pseudogenized *prfB* genes in the MGR-GC4 MAGs was established using sequence similarity search. Specifically, a protein database of *prfB* genes from 61 *Eggerthellaceae* genomes in GTDB R10-RS226 (1 per genus) was constructed and tBLASTn 2.12.0+ [85] used to compare these protein sequences to the nucleotide sequences of the MGR MAGs. This resulted in full length hits with ∼60% identity to the *prfB* genes in the 4 MGR-GC11 MAGs. By contrast, the 34 MGR-GC4 MAGs only had low sequence identity hits (∼40%) that corresponded to the *prfA* gene in these genomes.

A differential gene presence analysis was performed by considering all genes identified by Prokka with a gene name and counting the number of genomes in each group with this gene (**Supp. Table 5**). Results are given for MGR-GC4 MAGs, MGR-GC11 MAGs, and the 4 *Eggerthellaeae* genomes closest to the MGR clade (**Figure 1**). Identification of motility and cell envelope genes were determined by inspecting Prokka annotations for specified gene names (**Supp. Table 6**).

### Alignment of conserved protein

Protein families where UGA aligned to a conserved tryptophan residue were identified using TIGRFAMs 17.0 [86] and HMMER 3.4 [87], with identified TIGRFAM families aligned with hmmalign. Conserved tryptophan residues were identified across the 38 MGR MAGs and genomes from *Eggerthella lenta* (GCF_000024265.1), *Adlercreutzia equolifaciens* (GCF_000478885.), and *Phoenicibacter congonensis* (GCF_900169485.1). Alignments were screened to identify positions where a tryptophan residue was present in at least 90% of these 41 genomes and the five proteins most clearly illustrating the recoding of UGA to tryptophan manually identified: TIGR00002 (*rpsP*; ribosomal protein bS16), TIGR00088 (*trmD*; tRNA (guanine(37)-N(1))-methyltransferase), TIGR00633 (*xth*; exodeoxyribonuclease III), TIGR01034 (*metK*; methionine adenosyltransferase), and TIGR01514 (*pncB*; nicotinate phosphoribosyltransferase).

### Calculation of genomic properties

Genomic properties were determined for 250 *Eggerthellaceae* genomes (**Supp. Table 2**) and genomes from taxa that use genetic code 4 (**Table 2**). Specifically, genomes from GTDB R10-RS226 were considered and subjected to the same QC criteria as used for the MGR MAGs. Notably, many species that use genetic code 4 fail the GTDB QC criteria as they lack a sufficient number of phylogenetically marker genes due to their highly reduced genomes. Consequently, *Hodgkinia cicadicola, Nasuia deltocephalinicola, Stammera capleta*, and *Zinderia insecticola* genomes were identified using NCBI classifications. Genomes belonging to the order *Mycoplasmatales* were identified using the GTDB taxonomy and only genomes assembled from the type strain of a species retained to focus on high-quality genomes. The %GC content and genome size of these species were determined using CheckM2 and the presence of tRNAs and the *prfB* genes identified using Prokka. The frequency of stop codon usage was determined by parsing annotated CDS features and tabulating the number of identified stop codons.

### Gene neighbourhood of *secA* gene

Prokka annotations were used to identify *secA* genes (see *Identification of genes*) and a custom Python script used to create GFF files with the 20 genes adjacent on either side of the *sec*A gene. These GFF files were provided to Clinker 0.0.31 [88] to generate the gene neighbourhood plot which was enhanced with visual elements using Inkscape.

### Host and geographic metadata

Host and geographic metadata was collated for 967 SRA sample where SingleM [28] predicted the abundance of CAVGFB01 or JAUNQF01 at >0.1% (**Supp. Table 1**). Sample metadata was obtained from NCBI using the biosample2table.py script provided by the Jason Stajich Lab (https://github.com/stajichlab/biosample_metadata). The GTDB R10-RS226 SingleM database was supplemented with the 35 MGR MAGs and the 967 samples re-profiled to resolve the presence of taxa from the genera *Tapirivita, Equivita*, and *Gorillivita* proposed in this study (**Supp. Table 1**).

## Data availability

The 35 MGR MAGs were submitted to NCBI under BioProject PRJNA1366607. These MAGs were recovered from SRA samples (**Supp. Table 2**) and come from 13 BioProjects: PRJEB47977 (7 MAGs; unpublished), PRJEB73511 (1 MAG) [89], PRJNA1089803 (1 MAG) [89], PRJNA382701 (1 MAG) [90], PRJNA545604 (1 MAG; unpublished), PRJNA554776 (8 MAGs) [91], PRJNA635116 (3 MAGs) [92], PRJNA784453 (4 MAGs) [93], PRJNA814825 (2 MAGs) [94], PRJNA860652 (1 MAG) [95], PRJNA871798 (1 MAG; unpublished), PRJNA887424 (1 MAG) [49], and PRJNA983076 (4 MAGs) [31]. All unpublished data was submitted to NCBI at least 3 years prior to this manuscript. This manuscript adheres to the data reuse guidelines of Hug et al. [96].

## Acknowledgements

We gratefully acknowledge the significant effort expended in collecting, sequencing, and depositing metagenomic datasets into public repositories.

## Footnotes

1 https://www.insdc.org/submitting-standards/genetic-code-tables

