## Supplementary Information for "First report of stop codon reassignment to tryptophan in members of the bacterial phylum *Actinomycetota*" (from another version of this preprint: version 1, 2025-12-02)

#### Supplementary Notes

**Supplementary Note 1**. Here we provide analyses in support of dividing the JAUNQF01 MAGs into two genera. The primary rationale for defining two genera is to communicate that genomes in this group form two lineages that utilize different genetic codes. Division into two genera is supported by both relative evolutionary divergence (RED; Parks et al., 2018) and 16S rRNA percent identity (Hackmann, 2025).

RED values for *Eggerthellaceae* genera were assessed in a tree including the MGR MAGs and all GTDB R226 species representatives comprised of ≤200 contigs and with CheckM2 completeness ≥80% and contamination ≤5%. This tree was inferred using the GTDB R226 methodology (i.e. tree 1 in **Supp. Table 3**). RED values were determined using PhyloRank v0.1.12 (github.com/donovan-h-parks/PhyloRank). The mean RED value for the 39 *Eggerthellaceae* genera in GTDB R226 with genomes passing QC was 0.93 ± 0.035 (median = 0.93; minimum = 0.87; maximum = 0.99). The proposed taxa having a RED of 0.86 for *Gorillivita*, 0.89 for *Equivita*, and 0.93 for *Tapirivita*. These RED values being the same or lower than the mean RED value for *Eggerthellaceae* genera supports classifying these taxa as new *Eggerthellaceae* genera.

The 16S rRNA sequences in the *Tapirivita inops* (MGR6; 1517 bp) and *Equivita altericodex* (MGR7; 1514 bp) type genomes were compared to genomes in the NCBI core nucleotide database using BLAST v2.12.0 (Camacho et al., 2009). The *Tapirivita inops* 16S rRNA sequence had matches to a range of *Eggerthellaceae* sequences with the top hits being to *Raoultibacter timonensis* (87.7% identity; score = 1749) and *Phoenicibacter congonensis* (87.8% identity; score = 1720). Similarly, the *Equivita altericodex* 16S rRNA sequence had matches to a range of *Eggerthellaceae* species including *R. timonensis* (88.8% identity; score = 1847), *Eggerthella lenta* (88.6% identity; score = 1836), and *Phoenicibacter congonensis* (88.3% identity; score = 1799). These results are congruent with the phylogenetic position of these MGR species within the *Eggerthellaceae* family and indicate these 16S rRNA sequences are unlikely to represent contamination. The 16S rRNA identity between the *Tapirivita inops* and *Equivita altericodex* 16S rRNA sequences was determined to be 95.3% using the NCBI BLAST alignment tool (blast.ncbi.nlm.nih.gov; Zhang et al., 2000), a level of sequence similarity within the range observed for bacterial genera within the same family (95.9% being the 5th percentile; Hackmann, 2025). Notably, relatively high sequence similarity between 16S rRNA sequences from different genera within the *Eggerthellaceae* is not atypical, *e.g. Parvibacter caecicola* and *Adlercreutzia equolifaciens* 16S rRNA sequences have 96.1% identity, *Parvibacter caecicola* and *Enterorhabdus faecimuris* and 16S rRNA sequences have 96.0% identity, and *Enteroscipio rubneri* and *Arabiibacter massiliensis* 16S rRNA sequences have 95.2% identity.


Camacho C, et al. 2009. BLAST+: architecture and applications. *BMC Bioinformatics*, 10, 421.

Zheng Z, et al. 2000. A greedy algorithm for aligning DNA sequences, *J Comput Biol*, 7, 203-14.

### Supplementary Figures

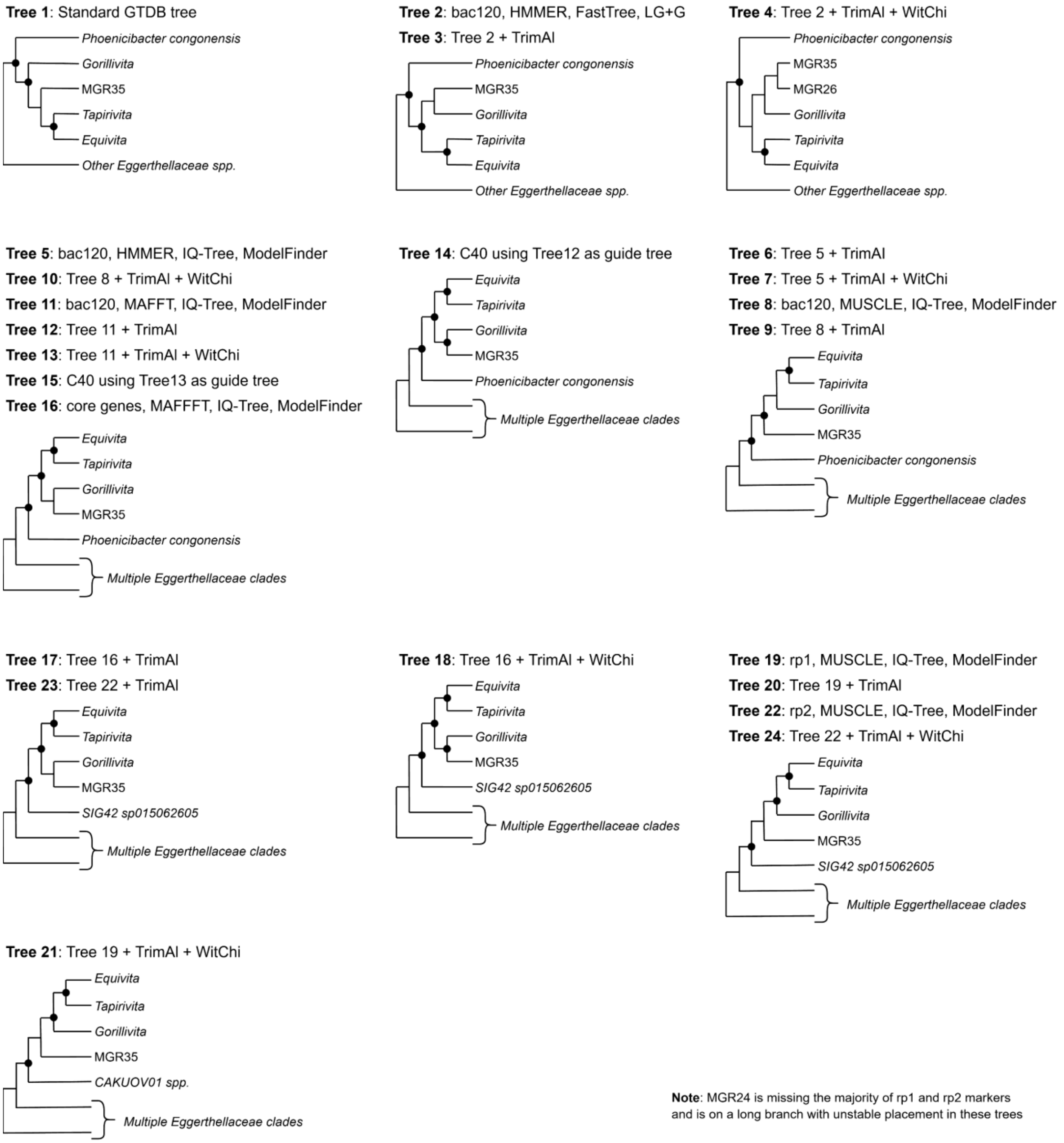


**Supp. Figure 1**. Topology of *Eggerthellaceae* species and recovered JAUNQF01, CAVGFB01, and MGR35 MAGs for 24 trees inferred using varying marker sets, MSA methodologies, inference methods, and evolutionary models (*see Methods* and **Supp. Table 3**).

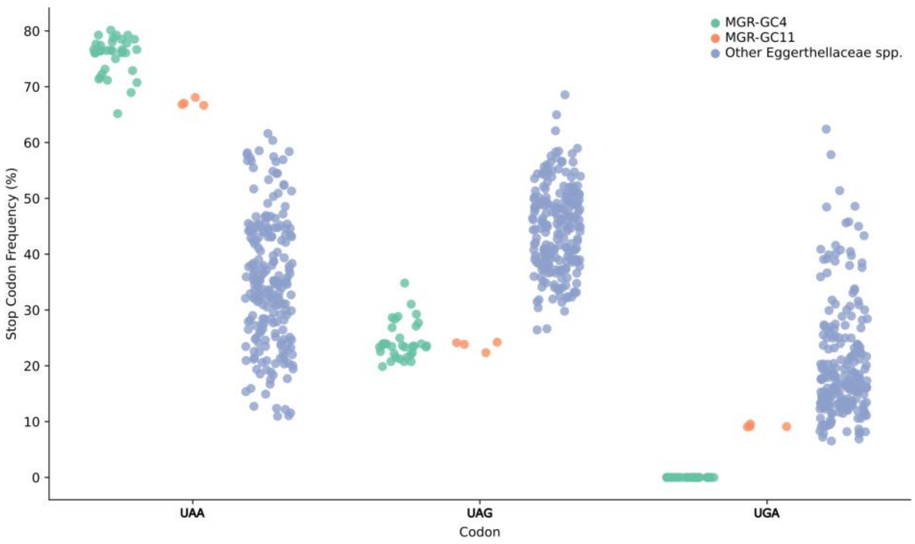


**Supp. Figure 2**. Frequency of stop codon usage for the 34 MGR-GC4 MAGs, 4 MGR-GC 11 MAGs, and 212 *Eggerthellaceae* genomes outside the MGR clade.

### Supplementary Tables

**Supp. Table 1**. Host range of samples predicted to contain JAUNQF01 and/or CAVGFB01 species (see Excel file).

**Supp. Table 2**. Metadata for recovered MGR MAGs and *Eggerthellaceae* genomes used in phylogenetic analyses (see Excel file).

**Supp. Table 3**. Description of novel taxa proposed in this study. SeqCode Register List: seqco.de/r:eiwpwrw3

| *Proposed taxon* | *Etymology* | *Nomenclatural type* | *Description* |
|---|---|---|---|
| *Tapirivita* gen. nov. | Ta.pi.ri.vi'ta. N.L. masc. n. *Tapirus* the genus name of the tapir; L. fem. n. *vita*, life; N.L. fem. n. *Tapirivita* a life form associated with a tapir | Species Tapirivita inops$^{Ts}$ | The description is identical to that given for the type species, *Tapirivita inops*. The genus was formerly known as JAUNQF01 in GTDB. Parent taxon: family *Eggerthellaceae*. |
| *Tapirivita inops* sp. nov. | in'ops. L. fem. adj. *inops*, helpless, destitute, indigent, poor, referring to its small genome size | Genomic assembly: XXX | The nomenclatural type, MAG MGR6, is a metagenome-assembled genome recovered from the gut microbiome of a tapir at Zoo Saarbruecken, Germany. The assembly was reconstructed from SRA reads generated in the study of Schmartz et al. (2024) (Biosample: SAMN35719516; BioProject: PRJNA983076). The assembly is of high quality with an estimated completeness of 93.05% and estimated contamination of 0.24% (CheckM v2) and encodes tRNAs for 20 amino acids. The genome size is 1.5 Mb, and contains full-length 23S, 16S and 5S rRNA genes. The GC content is 35.1%. Genomic characteristics suggest a host-dependent lifestyle in the mammalian gut, consistent with features of an obligate symbiont. |
| *Equivita* gen. nov. | E.qui.vi'ta. L. masc. n. *equus*, a horse; L. fem. n. *vita*, life; N.L. fem. n. *Equivita* a life form associated with a horse | Species *Equivita altericodex*$^{Ts}$ | The description is identical to that given for the type species, *Equivita altericodex*. The genus was formerly known as JAUNQF01 in GTDB. Parent taxon: family *Eggerthellaceae*. |
| *Equivita altericodex* sp. nov. | al.te.ri.co'dex. L. masc. pron. *alter,* another, the other; L. masc. n. *codex*, a book, code; N.L. masc. n. *altericodex* alternative code, referring to genetic recoding process | Genomic assembly: XXX | The nomenclatural type, MAG MGR7, is a metagenome-assembled genome recovered from the gut microbiome of a horse in Xinjiang, China. The assembly was reconstructed from SRA reads generated in the study of Zhou et al. (2022) (Biosample: SAMN23489792; BioProject: PRJNA784453). The assembly is of high quality with an estimated completeness of 90.46% and estimated contamination of 0.06% (CheckM v2) and encodes tRNAs for 20 amino acids. The genome size is 1.4 Mb, and contains a full-length 16S gene (23S and 5S rRNA genes are not detected). The GC content is 36.1%. Genomic characteristics suggest a host-dependent lifestyle in the mammalian gut. Genomic and phylogenetic evidence demonstrates that UGA encodes tryptophan in the genome of this species. It is hypothesized that host dependency may have facilitated this stop-to-tryptophan codon reassignment. |
| Gorillivita gen. nov. | Go.ril.li.vi'ta. N.L. masc. n. *Gorilla* the genus name of the great ape; L. fem. n. *vita*, life; N.L. fem. n. *Gorillivita* a life form associated with a gorilla | Species *Gorillivita intestinalis*$^{Ts}$ | The description is identical to that given for the type species, *Tapirivita inops*. The genus was formerly known as CAVGFB01 in GTDB. Parent taxon: family *Eggerthellaceae*. |
| *Gorillivita intestinalis* sp. nov. | in.tes.ti.na'lis. N.L. fem. adj. intestinalis, pertaining to the intestine | Genomic assembly: XXX | The nomenclatural type, MAG MGR2, is a metagenome-assembled genome recovered from the gut microbiome of a western lowland gorillas (*Gorilla gorilla gorilla*) at the Dzanga Sangha Protected Areas, Central African Republic. The assembly was reconstructed from SRA reads generated in the study of Sharma et al. (2020 )(Biosample: SAMN15047031; BioProject: PRJNA635116). The assembly is of high quality with an estimated completeness of 92.27% and estimated contamination of 0% (CheckM v2) and encodes tRNAs for 18 amino acids. The genome size is 0.9 Mb, with no detectable 16S, 23S and 5S rRNA genes. The GC content is 31.8%. Genomic characteristics suggest a host-dependent lifestyle in the mammalian gut. Genomic and phylogenetic evidence demonstrates that UGA encodes tryptophan in this genome. It is hypothesized that host dependency may have facilitated this stop-to-tryptophan codon reassignment. |

**Supp. Table 4**. Exploration of MGR monophyly, internal topology, and position relative to *Eggerthellaceae* genera for trees obtained using varying marker sets, alignment methods, alignment filtering methods, tree inference methods, and models of evolution.

| *Tree ID* | *Marker Set* | *Alignment Method* | *MSA Filtering (no. columns)* | *Tree Inference Method* |
|---|---|---|---|---|
| 1 | bac120 | HMMER | Standard GTDB filtering and random site selection (5,036) | FastTree with WAG (inferred over GTDB representative) |
| 2 | bac120 | HMMER | None (40,105) | FastTree with LG+G |
| 3 | bac120 | HMMER | TrimAl (37,029) | FastTree with LG+G |
| 4 | bac120 | HMMER | TrimAl + WitChi (28,621) | FastTree with LG+G |
| 5 | bac120 | HMMER | None (40,105) | IQ-Tree with Q.pfam+F+I+R10 |
| 6 | bac120 | HMMER | TrimAl (37,029) | IQ-Tree with LG+F+I+R10 |
| 7 | bac120 | HMMER | TrimAl + WitChi (28,621) | IQ-Tree with LG+F+R10 |
| 8 | bac120 | MUSCLE | None (89,321) | IQ-Tree with Q.pfam+F+I+R10 |
| 9 | bac120 | MUSCLE | TrimAl (43,657) | IQ-Tree with Q.pfam+F+I+R10 |
| 10 | bac120 | MUSCLE | TrimAl + WitChi (32,137) | IQ-Tree with LG+F+R10 |
| 11 | bac120 | MAFTT | None (42,597) | IQ-Tree with Q.pfam+F+R10 |
| 12 | bac120 | MAFTT | TrimAl (40,792) | IQ-Tree with Q.pfam+F+R10 |
| 13 | bac120 | MAFTT | TrimAl + WitChi (30,832) | IQ-Tree with LG+F+R10 |
| 14 | bac120 | MAFTT | TrimAl (40,792) | IQ-Tree with Q.pfam+F+C40 (Q.pfam+F+R10 as guide tree) |
| 15 | bac120 | MAFTT | TrimAl+ WitChi | IQ-Tree with LG+F+C40 (LG+F+R10 as guide tree) |
| 16 | core genes | MAFFT | None (172,238) | IQ-Tree with LG+F+I+R10 |
| 17 | core genes | MAFFT | TrimAl (108,222) | IQ-Tree with LG+F+I+R10 |
| 18 | core genes | MAFFT | TrimAl + WitChi (89,222) | IQ-Tree with LG+F+R10 |
| 19 | rp1 | MUSCLE | None (3,463) | IQ-Tree with Q.yeast+I+R7 |
| 20 | rp1 | MUSCLE | TrimAl (2,766) | IQ-Tree with Q.yeast+I+R7 |
| 21 | rp1 | MUSCLE | TrimAl + WitChi (2,606) | IQ-Tree with Q.yeast+I+R7 |
| 22 | rp2 | MUSCLE | None (5,833) | IQ-Tree with Q.yeast+R9 |
| 23 | rp2 | MUSCLE | TrimAl (4,065) | IQ-Tree with Q.yeast+I+R8 |
| 24 | rp2 | MUSCLE | TrimAl + WitChi (3,793) | IQ-Tree with Q.yeast+I+R7 |

**Supp. Table 5**. Differential gene presence analysis between MGR-GC4 MAGs, MGR-GC11 MAGs, and *Eggerthellaceae* genomes (see Excel file).

**Supp Table 6**. Presence of motility and cell envelope genes for MGR MAGs, *Eggerthellaceae* genomes, and select obligate host-dependent taxa (see Excel file).
